# Aversive learning strengthens episodic memory in both adolescents and adults

**DOI:** 10.1101/547331

**Authors:** Alexandra O. Cohen, Nicholas G. Matese, Anastasia Filimontseva, Xinxu Shen, Tracey C. Shi, Ethan Livne, Catherine A. Hartley

**Affiliations:** Department of Psychology, New York University; Columbia University Irving Medical Center; Weizmann Institute of Science

## Abstract

Adolescence is often filled with positive and negative emotional experiences that may change how individuals remember and respond to stimuli in their environment. In adults, aversive events can both enhance memory for associated stimuli as well as generalize to enhance memory for unreinforced but conceptually related stimuli. The present study tested whether learned aversive associations similarly lead to better memory and generalization across a category of stimuli in adolescents. Participants completed an olfactory Pavlovian category conditioning task in which trial-unique exemplars from one of two categories were partially reinforced with an aversive odor. Participants then returned 24-hours later to complete a recognition memory test. We found better corrected recognition memory for the reinforced versus the unreinforced category of stimuli in both adults and adolescents. Further analysis revealed that enhanced recognition memory was driven specifically by better memory for the reinforced exemplars. Autonomic arousal during learning was also related to subsequent memory. These findings build on previous work in adolescent and adult humans and rodents showing comparable acquisition of aversive Pavlovian conditioned responses across age groups and demonstrate that memory for stimuli with an acquired aversive association is enhanced in both adults and adolescents.

## Introduction

Emotional experiences shape the information that we remember. Emotional events, particularly those that are negative, have been widely shown to enhance episodic memory in human adults (Cahill & McGaugh, 1998; Labar & Cabeza, 2006; Yonelinas & Ritchey, 2015). However, studies examining whether emotion similarly facilitates episodic memory at earlier developmental stages have yielded mixed results. Studies of autobiographical memories for emotional and neutral events in children and adolescents suggest that emotional life events are remembered more frequently and in greater detail (Bauer et al., 2017; Fivush, Hazzard, McDermott Sales, Sarfati, & Brown, 2003). In contrast, several studies assessing children’s subsequent memory for images depicting intrinsically emotional stimuli have shown similar memory for negative and neutral images (Cordon, Melinder, Goodman, & Edelstein, 2013; Leventon, Stevens, & Bauer, 2014). There is some evidence to suggest that emotional information enhances memory in adolescents. Adolescents show enhanced memory for fearful faces relative to neutral faces (Pinabiaux et al., 2013) and their recall for emotional images is similar to that of adults (Vasa et al., 2011). In an attempt to reconcile these findings in more limited age-ranged samples, a recent study examined subsequent memory for negative, neutral, and positive images in 8- to 30-year-olds and showed similar emotional memory enhancement effects across ages (Stenson, Leventon, & Bauer, 2019). Taken, together, these studies suggest that across development, memory for emotional experiences may be better than memory for neutral experiences and that emotional memory facilitation may emerge relatively early in development, during childhood. Still, the extant research has focused on memory for events from one’s own life or for intrinsically emotional stimuli. Adolescence, in particular, is a stage of development associated with increased exploration and exposure to novel contexts, leading to many new and emotionally salient experiences (Casey, 2015). Thus, this may be a time when episodic memories for positive or negative associations are especially crucial for guiding future behaviors (Murty, Calabro, & Luna, 2016). Yet it remains unclear whether emotional learning, in which a neutral stimulus associated with an emotional experience acquires affective significance, similarly enhances subsequent memory in adolescents and adults.

Studies of emotional learning commonly model the acquisition of emotional associations through Pavlovian learning, in which a previously neutral conditioned stimulus acquires emotional salience through pairing with an intrinsically arousing positive or negative unconditioned stimulus (LeDoux, 2000; Maren, 2001). Although previous research suggests that acquisition of negative emotional associations is readily observable early in development (Deal, Erickson, Shiers, & Burman, 2016; Kim, Li, & Richardson, 2011; Pattwell et al., 2012; Rudy, 1993), changes in learning processes following acquisition (Baker, Bisby, & Richardson, 2016) suggest that there may be differences in the persistence of learned aversive associations in memory in adolescents relative to adults. Additionally, in real world situations, learned emotional associations are often more complex than an association between a simple stimulus and an emotionally salient outcome. For example, if someone is bitten by a dog, they may go on to develop a negative association not only with the particular dog that bit them, but with all dogs, or with animals more generally. The generalization and persistence in memory of learned aversive associations are core features of anxiety disorders (American Psychiatric Association, 2013; Dymond, Dunsmoor, Vervliet, Roche, & Hermans, 2015). Characterizing how these cognitive processes develop is particularly important given the typical emergence and peak in prevalence of anxiety disorders during adolescence (Kessler, Berglund, Demler, Jin, & Walters, 2005). In children and adolescents, stimuli that are visually similar to an aversive conditioned stimulus elicit more negative subjective emotion ratings and heightened psychophysiological measures of arousal, suggesting a generalization of negative affective value (Glenn et al., 2012; Michalska et al., 2016; Schiele et al., 2016). However, whether aversive learning generalizes more broadly to facilitate subsequent memory for similar stimuli in adolescence, a period of development in which anxiety disorders often first emerge, has yet to be investigated.

A recently developed “category conditioning” paradigm enables measurement of both learned affective responses and their generalization to conceptually similar stimuli, as well as the degree to which the strength and generalization of subsequent memory is influenced by emotionally salient events. In this paradigm, trial-unique exemplars from one conceptual category are partially reinforced with an intrinsically positive or negative stimulus, while exemplars from another conceptual category are never reinforced (Dunsmoor, Kragel, Martin, & La Bar, 2014; Dunsmoor, Martin, & LaBar, 2012; Patil, Murty, Dunsmoor, Phelps, & Davachi, 2017). In adults, emotional associations formed via category conditioning can generalize across the conceptual category and lead to enhanced memory for the reinforced category of exemplars (e.g. Dunsmoor et al., 2014, 2012; Dunsmoor, Murty, Davachi, & Phelps, 2015; Kroes, Dunsmoor, Lin, Evans, & Phelps, 2017; Patil et al., 2017). Here we leverage this paradigm to examine in both adults and adolescents whether emotional memory is facilitated for stimuli with an aversive association, whether such a memory benefit generalizes to non-reinforced exemplars within a category, and whether psychophysiological signatures of aversive learning also generalize to conceptually similar stimuli.

In the present study, 60 participants ages 13- to 25-years-old completed a novel olfactory Pavlovian category conditioning task, followed by a recognition memory test 24-hours later (Figure 1). Our category conditioning paradigm used aversive odor as an unconditioned stimulus, rather than mild electrical shock, as aversive odors have been successfully used in conditioning paradigms in human (Gottfried, O’Doherty, & Dolan, 2002) and non-human primates (U. Livneh & Paz, 2012; Uri Livneh & Paz, 2010, 2012) and can be ethically administered in developmental populations without risk of physical harm. We chose to administer the memory test a day after learning due to convergent evidence from previous studies suggesting that emotional memory enhancement effects emerge with time, after at least several hours (Yonelinas & Ritchey, 2015). The skin conductance response (SCR) to each stimulus was collected during the category conditioning task to serve as a psychophysiological measure of learning. Additionally, following the recognition memory test, we collected measures of self-reported anxiety and intolerance of uncertainty, which we hypothesized might relate to individual differences in emotional memory enhancement effects. Our primary aim was to test whether acquired aversive associations, using odor as a reinforcer, enhance memory in adolescents, similarly to adults, and whether these aversive associations generalize across a category. We hypothesized that adolescents and adults would show similar facilitation of memory for the aversively reinforced stimuli, but that adolescents might show greater generalization of aversive associations across a category relative to adults, which might confer heightened vulnerability to anxiety during this developmental stage.

**Figure 1.**
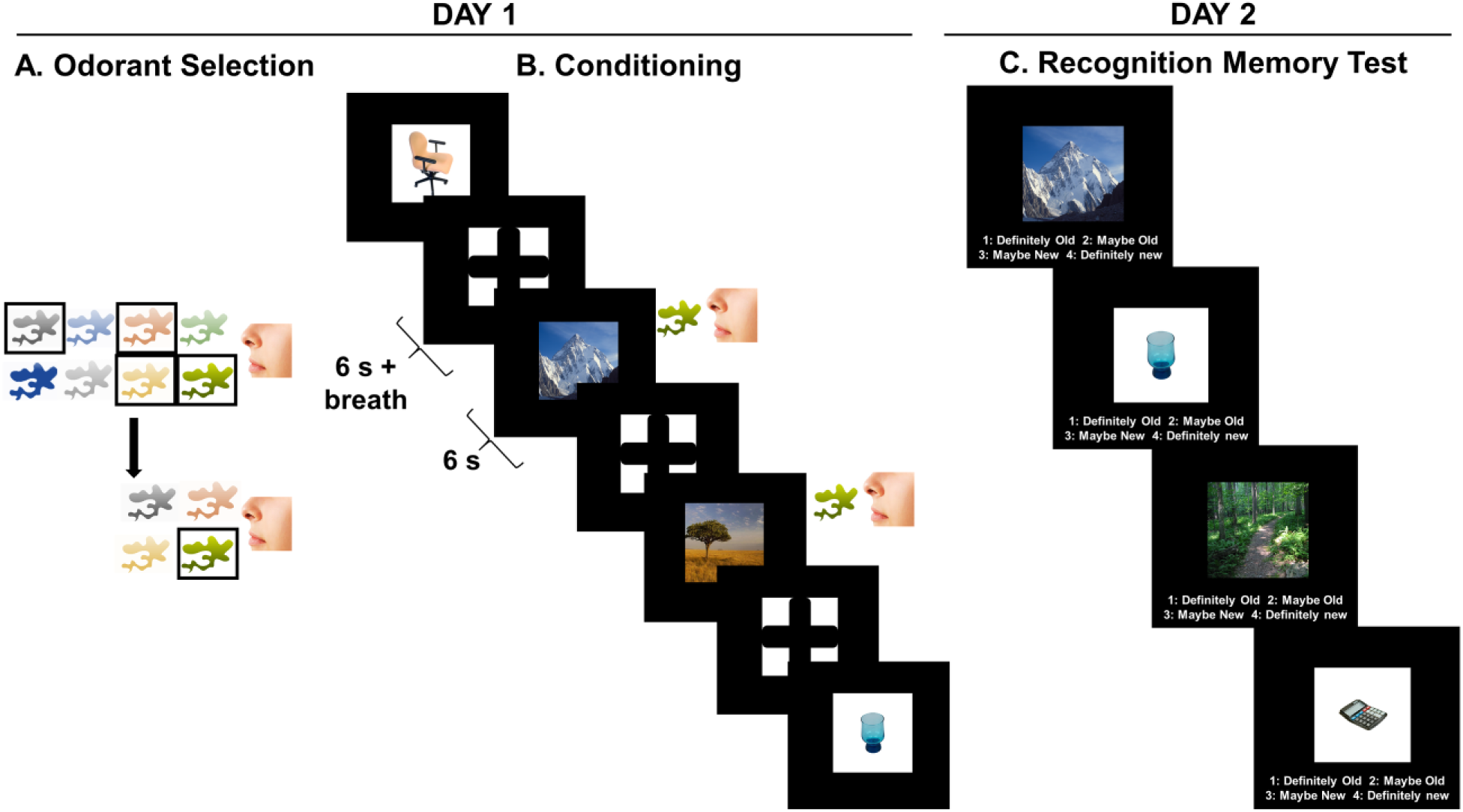
Experimental design. Participants first completed an odor selection procedure which involved a two-part rating procedure (A) to determine which odorant would be used as the unconditioned stimulus (US). Participants provided valence and arousal ratings for eight odorants. These ratings were used to select a set of four odorants that were delivered via the olfactometer and rated again to identify the final US (for more details see Methods). Immediately afterwards, participants underwent aversive olfactory Pavlovian category conditioning, using a breath-triggered paradigm, in which one category of images (CS+) was reinforced 50% of the time and the other category (CS-) were never reinforced (B). Participants returned 24-hours later and completed a self-paced recognition memory test that included all the images observed on day one, plus an equal number of new images from each category (C).

## Results

### Recognition Memory

In line with previous category conditioning studies (Dunsmoor et al., 2014, 2012, 2015), we first examined corrected recognition memory (hit minus false alarms) for stimuli from the reinforced (CS+) versus unreinforced (CS-) category by continuous age (Figure 2A) and controlling for which category (objects or places) served as the reinforced category. We found a significant effect of stimulus type (*F*(1,58) = 8.91, *p* = 0.004), such that subjects showed better corrected recognition memory for the CS+ stimuli than the CS-stimuli. There was no significant effect of age (*F*(1,57) = 0.31, *p* = 0.58), no age by stimulus type interaction (*F*(1,58) = 0.33, *p* = 0.57), and no effect of which category was reinforced (*F*(1,57) = 0.58, *p* = 0.45).

**Figure 2.**
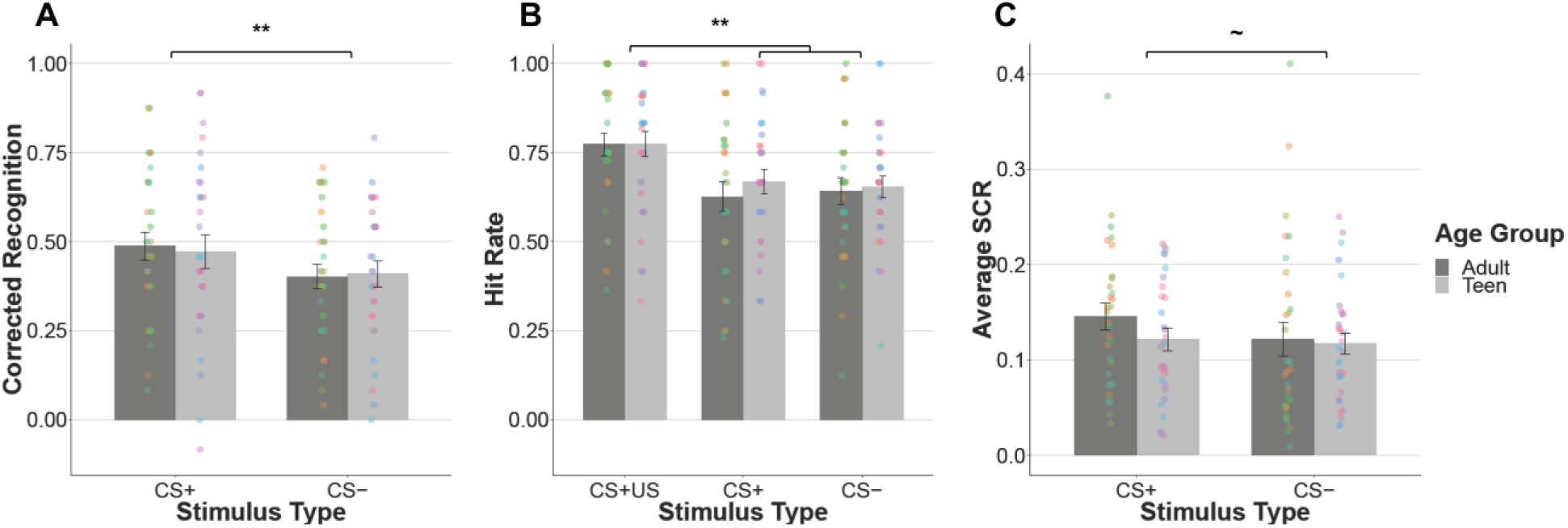
Similar effects of aversive learning on recognition memory and skin conductance response across age. Across age, corrected recognition memory is better for items from the CS+ versus CS-category (A), driven by better recognition memory for the reinforced items (CS+US) (B). There was a trend towards higher skin conductance in response to CS+ items relative to CS-items (C). Participants are separated by age group (Teen: 13-17, Adult: 18-25) for visualization purposes only. The corresponding statistical analyses treat age as a continuous variable. Different colored dots represent individual participants. Error bars are s.e.m. ** *p* < .01, ∼ *p* < .1

To assess whether the memory benefit conferred by learned aversive associations generalized to unreinforced exemplars within the same conceptual category, we next examined hit rate by stimulus type (CS+US, CS+, and CS-) and by continuous age (Figure 2B), controlling for the reinforced category. We found a significant effect of stimulus type (*F*(2,116) = 14.95, *p* < 0.0001), but no significant effects of age (*F*(1,57) = 0.69, *p* = 0.41), no age by stimulus type interaction (*F*(2,116) = 1.14, *p* = 0.32), and no effect of which category was reinforced (*F*(1,57) = 0.01, *p* = 0.93). Post-hoc t-tests (α = .0167, adjusted for multiple comparisons) revealed that the main effect of stimulus type was driven by better memory for the CS+US stimuli relative to the unreinforced CS+ stimuli (*t*(116.59) = 3.55, *p* < 0.001) and the CS-stimuli (*t*(117.99) = 3.73, *p* < 0.001). There was no significant difference between memory for the unreinforced CS+ stimuli relative to the CS-stimuli (*t*(116.79) = 0.02, *p* = 0.99). We also examined trial-wise memory accuracy (hit rate) by stimulus type (CS+US, CS+, and CS-) and by continuous age, controlling for both the reinforced category and the order of presentation of the stimuli during learning. We found a significant effect of stimulus type (*χ*^2^(2) = 42.38, *p* < .0001) and no significant effects of age (*χ*^2^(1) = 0.004, *p* = .95), no age by stimulus type interaction (*χ*^2^(2) = 3.82, *p* = .15), and no effect of which category was reinforced (*χ*^2^(1) = 0.22, *p* =.64). There was a significant primacy effect of stimulus presentation order (*χ*^2^(2) = 83.57, *p* < .0001), such that stimuli presented near the beginning of learning were better remembered than those presented near the end. These results suggest that improved corrected recognition memory for the CS+ category of stimuli was driven specifically by enhanced memory for images paired with an aversive odor and was not due to generalization of memory facilitation to non-reinforced images in the same category.

Given that several previous studies in adults observed such a generalization effect when analyzing the high confidence trials (e.g. Dunsmoor et al., 2012; Patil et al., 2017), we also conducted a post-hoc exploratory analysis in which we used an ordinal regression model (Burkner & Vuorre, 2018) to examine the influence of stimulus type on subsequent memory by confidence level. This approach allowed us to include all four confidence levels of memory responses (1 = Definitely Old, 2 = Maybe Old, 3 = Maybe New, and 4 = Definitely New). This analysis suggested that among the high-confidence hit stimuli, there is evidence for generalization of memory facilitation to unreinforced CS+ stimuli, such that there were more high confidence hits (1 = Definitely Old) than low confidence hits (2 = Maybe Old) for the both CS+US and CS+ stimuli, relative to the CS-stimuli, although the effect for CS+ stimuli is small. (see Supplemental Materials, Tables S1 & S2).

The pattern of results reported above remained consistent when participants who were excluded for not showing a variable skin-conductance signal during conditioning were included in the analyses (see Supplemental Materials, Figure S1).

### Psychophysiological Measure of Learning

To test for acquisition of category conditioning across adolescents and adults, we examined average skin conductance responses for stimuli from the reinforced (CS+) versus unreinforced (CS-) category by continuous age (Figure 2C). We found a marginal effect of stimulus type (*F*(1,58) = 3.03, *p* = 0.087), such that subjects showed a trend towards higher skin conductance for the CS+ stimuli relative to the CS-stimuli. There was no significant effect of age (*F*(1,58) = 1.50, *p* = 0.22) or an age by stimulus type interaction (*F*(1,58) = 0.60, *p* = 0.44). We also examined trial-by-trial unconditioned psychophysiological responses to trials that were paired with odors across the learning phase. We found a significant effect of trial number (*χ*2(1) = 31.94, p < .001), such that unconditioned responses decreased over the course of learning. This suggests that odor habituation did occur over the course of learning.

### Psychophysiology-Recognition Memory Relationships

To gain a better understanding of the large degree of individual variability in recognition memory, we explored relationships between psychophysiological responses during learning and subsequent memory. A number of previous studies in adults have shown that increased autonomic arousal during encoding is associated with better memory (Bradley, Greenwald, Petry, & Lang, 1992; Kensinger EA & Corkin S, 2004; Kleinsmith & Kaplan, 1963), therefore we first examined the relationship between subjects’ averaged skin conductance in response to all CS presentations across the encoding task and related this to overall memory performance (hit rate). We found a positive relationship (*r*(58) = 0.26, *p* = 0.041), such that individuals with larger conditioned responses showed better memory overall (Figure 3A).

**Figure 3.**
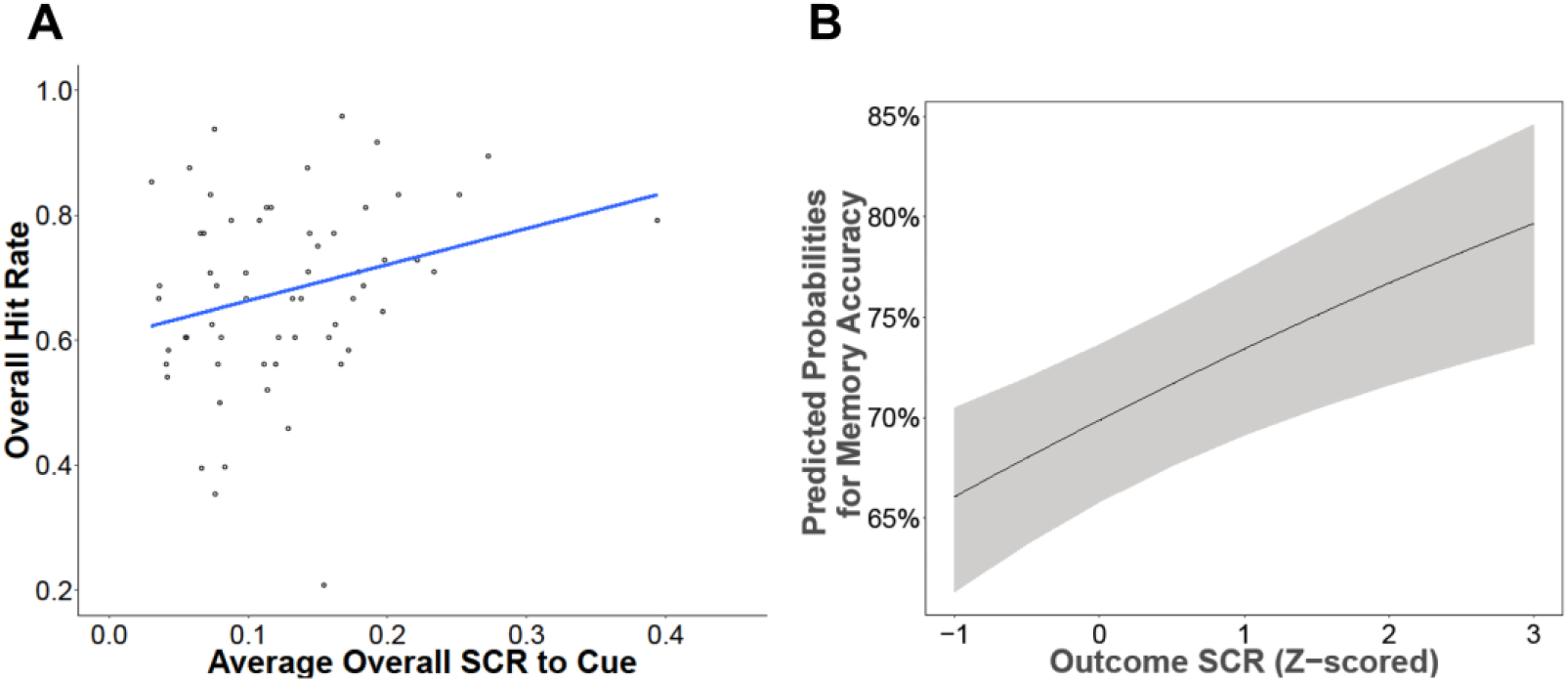
Psychophysiological arousal during learning relates to memory 24-hours later. Participants’ average skin conductance response to cue presentation was positively correlated with their overall recognition memory performance (A). While trial-evoked responses to the cue did not predict subsequent memory for that item, higher responses at the time of the outcome were predictive of better item memory (B).

We next investigated the relationship between trial-evoked skin conductance responses to CS presentation, irrespective of the stimulus type and experienced outcome, and subsequent memory. In the present study, we examined skin conductance responses at two time-points during each trial. The first time-point was when the CS was on the screen before the event occurred, which we refer to as responses to the cue. The second time-point began at the onset of the olfactometer “shoot” event (the release of either the odor or clean air), which we refer to as responses to the outcome. We first examined whether trial-evoked psychophysiological responses to the cue predicted subsequent memory, including continuous age as a regressor of interest. We found no significant effects of skin conductance in response to the cue (*χ*^2^(1) = 0.013, *p* = .91), age (*χ*^2^(1) = 0.34, *p* = .56), and no interaction between skin conductance in response to the cue and age (*χ*^2^(1) = 0.003, *p* = .95). We also examined whether trial-evoked skin conductance responses at the time of the outcome predicted subsequent memory, including continuous age as a regressor of interest. We found a significant main effect (*χ*^2^(1) = 14.11, *p* < .001), such that larger skin conductance responses at the time of the outcome were associated with better memory (Figure 3C). There was no statistically significant effect of age (*χ*^2^(1) = 0.22, *p* = .64), or interaction between outcome SCR and age (*χ*^2^(1) = 0.081, *p* = .77). Thus, trial-evoked SCRs at the time of outcome were predictive of subsequent memory whereas trial-evoked SCRs to the cue itself were not.

### Recognition Memory-Individual Difference Measure Relationships

To examine how individual differences in memory enhancement and generalization might relate to participants’ state or trait anxiety, and intolerance of uncertainty, we first computed memory bias scores for CS+US (CS+US hit rate – CS-hit rate) and unreinforced CS+ (CS+ hit rate – CS-hit rate) stimuli. Consistent with our earlier reported findings, linear regressions revealed no relationships between CS+US memory bias and age (*F*(1,58) = 0.18, *p* = 0.67) or CS+ memory bias and age (*F*(1,58) = 1.78, *p* = 0.19).

We next examined the relationships between the State Trait Anxiety Inventory (STAI) state and trait scales (Spielberger, Gorsuch, & Lushene, 1988) and these memory bias measures (α = .0125, adjusted for multiple comparisons). Linear regressions revealed no significant relationships between age and STAI state (*F*(1,58) = 0.51, *p* = 0.48), STAI trait (*F*(1,57) = 1.40, *p* = 0.24), or IUS (*F*(1,58) = 0.74, *p* = 0.38). We did not find statistically significant relationships between the STAI state measure and either memory bias index (CS+US memory bias, *r*(58) = -0.050, *p* = 0.71; CS+ memory bias, *r*(58) = -0.084, *p* = 0.52) or the STAI trait and CS+US memory bias (*r*(57) = -0.19, *p* = 0.15). However, we did observe a negative correlation between the STAI trait and CS+ memory bias (*r*(57) = -0.35, *p* = 0.006), such that individuals with lower STAI trait scores showed a stronger CS+ memory bias than those with high STAI trait scores. A follow-up analysis (α = .017, adjusted for multiple comparisons) was conducted to determine whether this result was due to differences in recognition memory for the unreinforced CS+ stimuli or the CS-stimuli. We also examined the relationship between recognition memory for the CS+US stimuli and STAI trait for completeness. We found that neither CS+US hit rate (*r*(57) = 0.09, *p* = 0.48) nor CS+ hit rate (*r*(57) = -0.11, *p* = 0.40) correlated with STAI trait scores. However, we observed a positive correlation between CS-hit rate and STAI (*r*(57) = 0.31, *p* = 0.016), such that individuals with higher trait anxiety showed better memory for the CS-stimuli (Figure 4A). A follow-up multiple-regression analysis including a STAI by continuous age interaction term revealed a significant main effect of trait anxiety (*F*(1,55) = 6.30, *p* = 0.015), no main effect of age (*F*(1,55) = 0.18, *p* = 0.675), and a marginal trait anxiety by age interaction (*F*(1,55) = 2.85, *p* = 0.097), such that the relationship between trait anxiety and memory for CS-stimuli was stronger in younger participants. These results indicate that higher self-reported anxiety was related to better for stimuli from the non-reinforced category, but was not related to memory for stimuli from the reinforced category.

**Figure 4.**
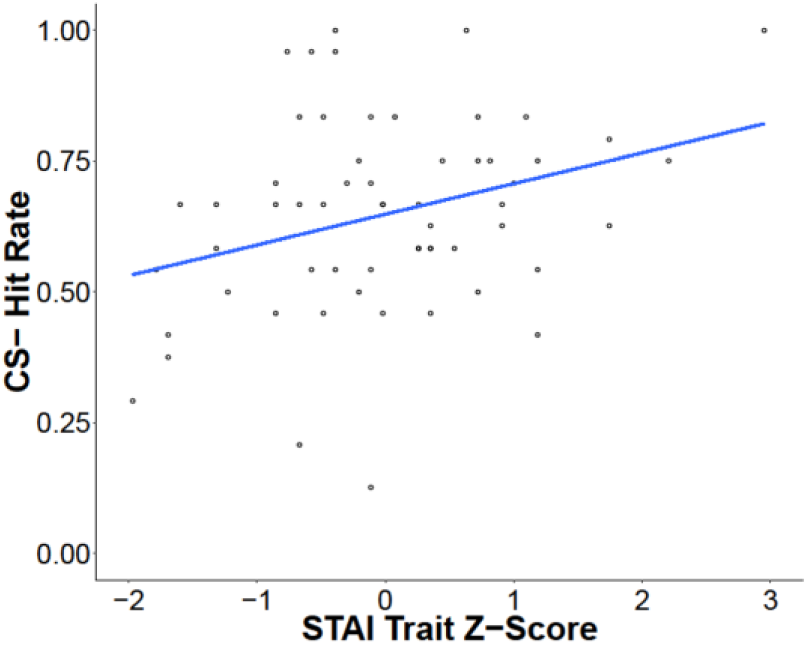
Better memory for the stimuli from the unreinforced category is associated with higher trait anxiety. While there was no significant relationship between recognition memory for the CS+ or CS+US stimuli and trait anxiety, there was a positive relationship between memory for the CS-stimuli and trait anxiety.

We also examined the relationships between Intolerance of Uncertainty Scale (IUS) (Freeston, Rhéaume, Letarte, Dugas, & Ladouceur, 1994) and the memory bias measures (α = .025, adjusted for multiple comparisons). We did not observe statistically significant correlations between either memory bias index and IUS (CS+US memory bias, *r*(58) = -0.053, *p* = 0.69; CS+ bias, *r*(58) = -0.21, *p* = 0.11).

The pattern of results reported above remained consistent when participants who were excluded for not showing a variable skin-conductance signal during conditioning were included in the analyses (see Supplemental Materials, Figure S2).

## Discussion

The present study employed a novel olfactory variant of a Pavlovian category conditioning task to test whether aversive learning leads to similar memory enhancement and generalization across a conceptual category in adults and adolescents. By using trial-unique stimuli as “tags” for each learning trial, we show that aversive learning leads to better episodic memory for trials associated with an aversive event in both adolescents and adults. The age invariance of this effect is consistent with previous observations that adolescent and adult humans and rodents exhibit equivalent acquisition of aversive Pavlovian conditioning using simple stimuli (Deal et al., 2016; Kim et al., 2011; Pattwell et al., 2012; Rudy, 1993). Our finding extends this literature by testing memory for individual events during conditioning and showing similar memory improvements in adolescents and adults for items with an acquired aversive association.

While few studies have examined the neural mechanisms underlying emotional facilitation of episodic memory prior to adulthood, our findings are consistent with evidence of the early development of this circuitry. Multiple memory systems, centered on the amygdala for emotional memory and the hippocampus for episodic and declarative memory, are proposed to interact to facilitate memory of emotional events (Mcdonald, Devan, & Hong, 2004; Phelps, 2004). Under the “emotional binding” account of episodic memory, the amygdala binds emotional information to an item, communicating with both the perirhinal cortex and the hippocampus to modulate encoding, storage, and recollection of these memories (Yonelinas & Ritchey, 2015). Evidence from rodent studies suggests that signatures of a functional emotional memory system emerge early in development (Stanton, 2000), indicating that emotional memory enhancement effects should be present during childhood and adolescence.

While memory for items directly associated with an aversive odor was facilitated across age, unreinforced exemplars from the same category as the odor-paired stimuli were not better remembered in either adults or adolescents. This result does not fully replicate previous studies showing that emotional associations generalize across a category and lead to enhanced memory for the reinforced category of exemplars in adults (Dunsmoor et al., 2014, 2012, 2015; Kroes et al., 2017; Patil et al., 2017). However, we did find some evidence of increased correct high-confidence memory judgements for both odor-paired stimuli and unreinforced stimuli of the same category, relative to stimuli that were never reinforced. This indicates a possible interaction between metacognitive ability and memory generalization effects, such that generalization of memory facilitation for unreinforced items from the same conceptual category as those that were aversively reinforced is primarily observed when examining high confidence memories. Alternatively, because the present study did not include a “don’t know” response option, generalization of emotional associations in memory may be obscured by noisiness in low confidence memory judgments due to guessing responses.

There were several differences between the present paradigm and the category conditioning paradigm used in previous work that may have contributed the lack of generalization of memory facilitation. The present paradigm used trial-unique object and scene images rather than tool and animal images. Objects were used to try and ensure that younger participants would have familiarity with the images and scenes were used instead of animals due to pilot data that suggested a general memory advantage for animals. It is possible that the exemplars from each category were too distinct to allow for generalization (Dunsmoor & Murphy, 2015). We also did not include expectancy ratings during conditioning in order to mitigate potential effects of generating a prediction on learning (Brod, Hasselhorn, & Bunge, 2018) and effects of expectancy rating on skin conductance response (Atlas et al., 2015). Although other variants of category condition paradigms have also omitted expectancy ratings and still observed memory facilitation effects (Patil et al., 2017), it is possible that this may have reduced the demand on participants’ attention, attenuating their anticipatory responses. Another reason that we may not have fully replicated prior studies is our use of a different primary reinforcer. Previous work has shown that the intensity of the aversive stimulus is related to the degree of generalization (Dunsmoor, Kroes, Braren, & Phelps, 2017), suggesting that olfactory reinforcers may not be potent enough to induce widespread generalization effects. We also saw evidence for habituation of the skin conductance response to the odor after repeated exposures across learning, which may have contributed in part to the observed primacy effect on memory. Further studies comparing aversive learning across different modality reinforcers (e.g. shock versus noise versus odor) and manipulating the duration and intensity of reinforcement will be necessary to determine the effectiveness of odor conditioning in producing generalization effects.

In this study, we used cue-evoked skin conductance response as a psychophysiological measure of emotional learning. Moreover, in this category condition paradigm, the measure of anticipatory arousal also provides a measure of the degree to which learned aversive associations generalize across a conceptual category. Skin conductance responses showed a trend towards increased anticipatory arousal for the reinforced category of stimuli across participants. While this marginal increase in anticipatory arousal indicates some degree of learning of the association between the partially reinforced category and a potential aversive outcome, evidence for emotional learning in our study was weak. In the current experiment, we used skin conductance response as a psychophysiological index of learning due to the prevalence of this measure in the human conditioning literature (Bradley, Miccoli, Escrig, & Lang, 2008; Hamm & Stark, 1993; LaBar, Gatenby, Gore, LeDoux, & Phelps, 1998; LANG, GREENWALD, BRADLEY, & HAMM, 1993). Other psychophysiological measures of learning, such as pupillometry (Leuchs, Schneider, & Spoormaker, 2018) and breathing measures (Uri Livneh & Paz, 2010) should also be examined to determine whether they might provide more robust indices of learning dynamics during olfactory conditioning.

In order to probe individual variability in aversive learning and memory, we examined relationships between skin conductance responses during learning and subsequent memory. Previous studies in adults have shown that autonomic arousal during encoding is associated with better memory (Bradley et al., 1992; Kensinger EA & Corkin S, 2004; Kleinsmith & Kaplan, 1963). In both humans and rodents (Glascher, Adolphs, Gläscher, & Adolphs, 2003; Mather, Clewett, Sakaki, & Harley, 2016; Reis & LeDoux, 1987; Reyes, Carvalho, Vakharia, & Van Bockstaele, 2011; Roozendaal, Luyten, de Voogd, & Hermans, 2016), the amygdala can modulate noradrenergic autonomic arousal responses to aversive stimuli, providing a putative mechanism through which emotion might influence memory. Consistent with this prior work, we found that individuals showing higher anticipatory arousal in response to cues on average, throughout the task, also showed better memory overall. However, in accordance with previous findings (de Voogd et al., 2016), trial-evoked anticipatory arousal to the cue did not predict subsequent memory for the corresponding trial. We instead found that trial-evoked responses at the time of the outcome predicted memory 24-hours later. These data suggest that while increased anticipatory arousal during learning may foster a general memory benefit, unlearned autonomic arousal reactions to individual stimuli predict whether or not that stimulus will be remembered at a later time.

Finally, we examined how individual difference measures related to subsequent memory. Unexpectedly, we found a positive relationship between recognition memory for CS-stimuli and trait anxiety, such that individuals with higher trait anxiety showed better memory for items from the category that was never reinforced. A follow-up analysis suggested that this correlation was largely driven by adolescent participants, although the trait anxiety by age interaction was only significant at a trend level. This result indicates that self-reported anxiety may promote memory for “safe” stimuli, which were never previously associated with an aversive outcome, within a context where aversive outcomes were experienced. While unexpected, this finding is consistent with the idea that overgeneralization of aversive experiences to dissimilar stimuli is a defining feature of anxiety (Dymond et al., 2015). In overgeneralization, the heightened emotional responses elicited by a threat-predictive stimulus are also displayed in response to other increasingly dissimilar stimuli. Our observation that memory for safe stimuli is facilitated in subjects with higher trait anxiety suggests that the extent to which the cognitive processes evoked by aversive experiences generalize to safe stimuli is also heightened in high anxiety individuals. These results also suggest that a relationship between anxiety and better memory for safe stimuli experienced within an aversive context may be more readily observable during adolescence, the period of development in which anxiety disorders often first emerge (Kessler et al., 2005; Lee et al., 2014). However, it is noteworthy that trait anxiety was not correlated with memory for the specific items associated with aversive events (CS+US stimuli) or the unreinforced items from that same category (CS+ stimuli). Given the exploratory nature of these results, replication and further investigation of the relationship between generalization, overgeneralization, and trait anxiety in adolescents and adults is warranted.

In summary, the current study demonstrated that aversive learning enhances episodic memory in both adolescents and adults, particularly for items directly associated with an aversive odor. We found that autonomic arousal during learning was related to later memory. Specifically, unlearned arousal responses to outcomes during encoding were predictive of subsequent memory for individual stimuli. These results indicate that aversive odors are sufficiently evocative to induce memory enhancements in both adolescents and adults. While further refinement of olfactory conditioning methods for use in developmental populations is necessary, this study suggests that aversive and appetitive odors might be fruitfully utilized to study emotional learning and memory processes across development.

## Materials and Methods

### Participants

Sixty participants between the ages of 13 and 25 years (mean age = 18.69, 30 female) were included in analyses. A target sample size of n = 60, including 30 adolescents and 30 adults, was determined based on group sample sizes previously reported in category conditioning studies (Dunsmoor et al., 2014, 2012, 2015). Data from 28 additional participants (mean age = 19.45, 18 female) were excluded from primary analyses for not showing a variable skin-conductance signal (defined as fewer than four scorable trials) during conditioning. Data from 22 additional participants were excluded from analyses due to the discovery of a software bug that yielded inconsistencies in timing and delivery of the aversive reinforcers. Three additional participants were excluded from analyses due to failure to return for the second session of the study. All participants were volunteers from a community sample of New York City. Of the 60 participants included in primary analyses, 45% of participants self-identified as Caucasian/White, 15% as African American, 25% as Asian, and 15% as mixed race. Additionally, 16.67% of the sample identified as Hispanic. Of the 28 participants who did not exhibit a variable skin conductance signal (but were included in the supplemental analyses of the memory data), 32.14% of participants self-identified as Caucasian/White, 21.43% as African American, 39.29% as Asian, and 7.14% as mixed race. Additionally, 7.14% of the sample identified as Hispanic. Participants were screened for difficulties seeing without corrective lenses (as the nasal mask precluded the simultaneous use of glasses; contact lenses were permitted), history of psychiatric diagnoses, use of psychoactive medication or beta blockers, or difficulties breathing. Participants provided informed written consent (adults) or assent (minors) per research procedures approved by New York University’s Institutional Review Board. Parents or guardians of teenagers under age eighteen also provided written consent on behalf of the teenager, prior to their participation in the study. All participants were compensated $30 for their participation in two approximately 1-hour sessions scheduled 24-hours apart.

### Olfactory Pavlovian Category Conditioning Paradigm

A category conditioning paradigm used in previous studies in healthy adults (Dunsmoor et al., 2014, 2012) was adapted for use with a custom built olfactometer, allowing for odorants to serve as the unconditioned stimulus (US). The breath-triggered conditioning paradigm consisted of four 12-trial blocks, where each trial was a unique exemplar from one of two conceptual categories. Over the course of conditioning, subjects viewed 24 unique objects and 24 unique scenes (Konkle & Caramazza, 2013). Each stimulus category (object or scene) was randomly assigned to serve as the reinforced conditioned stimulus category (CS+) for half the participants and the unreinforced conditioned stimulus category (CS-) for the other half of participants. Within each 12-trial block, half of the trials were exemplars from the CS+ category and half were from the CS-category. The CS-trials were never paired with an odor and the CS+ trials were reinforced 50% of the time (three CS+US, three CS+, and six CS-trials per block, resulting in 12 trials total). Trial order was pseudorandomized such that no more than two reinforced (CS+US) trials and no more than three exemplars from the same category appeared in a row. Four different trial orders and two possible assignments the reinforced category of stimuli resulted in eight different versions of the task that were administered to participants.

During conditioning, participants passively viewed images presented on the screen via Psychtoolbox-3 in Matlab R2015b while breathing through a nasal mask connected to the olfactometer. They were instructed simply to notice any associations between the pictures and smells. Clean air was continuously circulated through the mask and participants’ nasal breathing was measured via pressure sensors in the olfactometer and processed in real-time using LabVIEW 2016 Version 16.0f5 (64-bit). The paradigm used breath-triggered stimulus presentation to ensure that odor delivery was timed to a participants’ inhalation. Each trial began with a fixation cross presented for a fixed interval of six seconds. On the subsequent inhale after the fixation (variable duration), a trial-unique exemplar appeared on the screen for a fixed interval of six seconds. After a one-second buffer to ensure separation of respiratory cycles (Uri Livneh & Paz, 2010), the participant’s next inhale while the image was still on the screen triggered an olfactometer “shoot” event for two seconds. For CS- and unreinforced CS+ trials, this shoot event consisted of continued release of clean air and for the CS+US trials, the aversive odor was delivered. If the participant did not inhale while the image was on the screen, an olfactometer shoot event was not triggered and the participant only experienced clean air. If this occurred on a CS+US trial, this trial was reclassified as a CS+ image without reinforcement during data processing and in subsequent analyses. Twelve of the 60 participants missed at least one odor shoot event. Nine of the 12 missed a single shoot event, leading to a 45.83% reinforcement rate, two missed two shoot events, leading to a 41.67% reinforcement rate, and one participant missed three shoot events, leading to a 37.5% reinforcement rate.

### Odorant Selection

At the beginning of the first session, participants underwent an odor selection procedure to identify the odorant to be used as the aversive reinforcer in the category conditioning paradigm. This procedure was designed to take into account individual differences in whether an odorant is considered to be aversive, mirroring calibration procedures that are typically performed in aversive learning studies using mild electrical shock as the aversive reinforcer (e.g. Dunsmoor et al., 2014; Dunsmoor, Mitroff, & LaBar, 2009). Each odorant was rated on valence and arousal using a modified version of the Self-Assessment Manikin (SAM) (Bradley, M. M. & Lang, P. J., 1994). A suite of eight different aversive odorants, supplied by DreamAir perfumers, were first administered to participants using Whispi air puff canisters (Scentovation, Novia Products, LLC). Three of the odorants were the following chemical compounds: isovaleric aldehyde 10% diluted in isopropyl myristate, dimethyl acetate undecadienol, and Ozonil™ (tridec2-ene nitrile). Five of the odorants were proprietary DreamAir odorant blends (“Bad smell 3”, “Horse hair”, “Fumier”, “Frog 3”, and “Fear 45l”). Participants were asked to rate the valence of the smell on a scale from one to nine in which a one represented a bad smell, labeled ‘Don’t Like’ on the scale, and a nine represented a good smell, labeled ‘Like’ on the scale. Immediately following the valence rating, the arousal rating measured the perceived strength of the smell on a scale of one to nine in which one represented a ‘Weak’, unnoticeable odor and a nine represented a ‘Strong’, noticeable odor.

Each smell was presented and ranked on these scales three times, and the average scores for each odorant were computed to determine the four most aversive odors, as indexed by ratings of lowest valence and highest strength. The four most aversive odors were then presented through the nasal mask via inhale-triggered odor release delivered using the olfactometer, which allowed the participants to experience the odors as they would during conditioning. Participants were asked to rate the four odorants three times each on a scale from one to five, where one indicated the smell was bad and five indicated that the smell was so bad that the participant would not be able to handle smelling it several times during the conditioning task. Ratings were averaged and the odorant with the highest average of a score of four or below, meaning that the odor never received a rating of five, was used in the conditioning task.

### Recognition Memory Test

Participants returned 24-hours later for a recognition memory test presented via MATLAB’s Psychtoolbox-3. Participants were not told about the memory test until they arrived for the second session, at which point they were queried about their expectations for the session. Though the majority of participants reported no expectations, four of the thirty adults and two of the thirty teens reported that they anticipated some form of memory test. The self-paced memory test included the 24 CS+ and 24 CS-category exemplars from day one, as well as 24 new objects and 24 new scenes, for a total of 96 images. Images used in the task on day 1 and as new images on day 2 were counterbalanced across participants. Participants rated whether each picture was new or old on a four-point scale: 1 = Definitely Old, 2 = Maybe Old, 3 = Maybe New, and 4 = Definitely New. Consistent with previous studies, responses were collapsed across new versus old ratings. We examined corrected recognition memory, which is a difference score between hits, old images correctly identified as old, and false alarms, new images incorrectly identified as old. Additionally, we examined hit rate for the CS+US, CS+, and CS-images to look for generalization across the reinforced category of exemplars.

### Psychophysiological Data Acquisition & Analysis

Skin conductance data was recording during the conditioning paradigm using a BIOPAC MP-100 System (Goleta, CA). Pre-gelled SCR electrodes were placed on the hypothenar eminence of the palm (Dunsmoor et al., 2015) of the non-dominant hand and the phasic skin conductance response (SCR) to each CS onset and outcome timepoint (US or no US) were scored using AcqKnowledge 3.9 software (BIOPAC Systems). SCR data were low-pass filtered and smoothed. SCR scores were based on the window 0.5 seconds after stimulus onset to 0.5 seconds after shoot onset and outcome response scores were based on the window 0.5 seconds after shoot onset to 0.5 seconds after shoot offset. The trough-to-peak difference of the first waveform (in μSiem) (Dunsmoor et al., 2015; Hermans et al., 2017) beginning within these windows was measured. Using MATLAB R2016a, distributions were normalized using square root transformation of the raw SCR magnitudes, and then divided by the maximum response (across all cue and outcome responses) to enable between-subject comparison. Any trial without a shoot event was considered missing for analyses that examined SCR at outcome.

### Self-Report Measures

Following the recognition memory test on day two, participants completed several self-report measures via Qualtrics surveys. Participants completed the State Trait Anxiety Inventory (STAI) state and trait scales (Spielberger et al., 1988), the Intolerance of Uncertainty Scale (IUS) for adults (Freeston et al., 1994) or the IUS-C for teenagers ages 13 to 17 (Comer et al., 2010), and a free response question asking whether the subject noticed anything about the types of images that were paired with smells. One adolescent participant (16.96 year-old male) did not complete the STAI trait scale.

### Analysis Approach

Data processing was completed in MATLAB R2016 and all statistical analyses were conducted in R version 3.5.1 (R Core team, 2016). Mixed-effects models were run using the ‘lme4’ package (version 1.1-17) lmer (for analyzing recognition memory and average skin conductance response) and glmer (for trial-wise analyses) functions (Bates D, Maechler M, Bolker B, & Walker S, 2015). Numeric variables included as regressors in the model were z-scored across all participants. Each model included a random intercept for each participant. Statistics were reported from analysis of variance (Type III using Satterthwaite’s method) performed on lmer models and analysis of deviance (Type III Wald chi-square tests) performed on glmer models. Welch two sample t-tests were performed for post-hoc analyses of recognition memory data and Pearson product-moment correlations were computed for all reported correlations. Where applicable, statistical significance thresholds (alphas) adjusted for multiple comparisons are reported in the Results section.

### Data and code availability

Data and code are available on Open Science Framework: https://osf.io/qcx8t/

## Acknowledgements

We thank DreamAir, LLC for providing us with the odorants used in this study, Noam Sobel and his engineering team for building our olfactometer, and the Frueauff Foundation for a generous equipment grant that funded its construction. This work was supported by a Klingenstein-Simons Fellowship in Neuroscience, a Jacobs Foundation Research Fellowship, a NARSAD Young Investigator Award, the NYU Vulnerable Brain Project, and a National Science Foundation CAREER Award Grant No. 1654393 (to C.A.H.) as well as a National Science Foundation SBE Postdoctoral Research Fellowship Grant No. 1714321 (to A.O.C).

