## Supplement for "Aversive learning strengthens episodic memory in both adolescents and adults" (from another version of this preprint: version 2, 2019-02-13)

### Supplemental Material

We repeated all analyses of recognition memory reported in the main text including the 28 subjects that were excluded from analysis in the main text due to the lack of a variable skin-conductance signal. All results remained consistent with those reported in the main text. The results reported below include 88 participants between the ages of 13 and 25 years (mean age = 18.93, 49 female).

*Recognition Memory*

We examined corrected recognition memory and found a significant effect of stimulus type ($F(1,86) = 15.05$, $p < 0.001$), such that subjects showed better corrected recognition memory for the CS+ stimuli than the CS- stimuli (Figure S1A). There was not a significant effect of age ($F(1,85) = 0.13$, $p = 0.72$), an age by stimulus type interaction ($F(1,86) = 0.13$, $p = 0.72$), or an effect of which category was reinforced ($F(1,85) = 0.09$, $p = 0.77$).

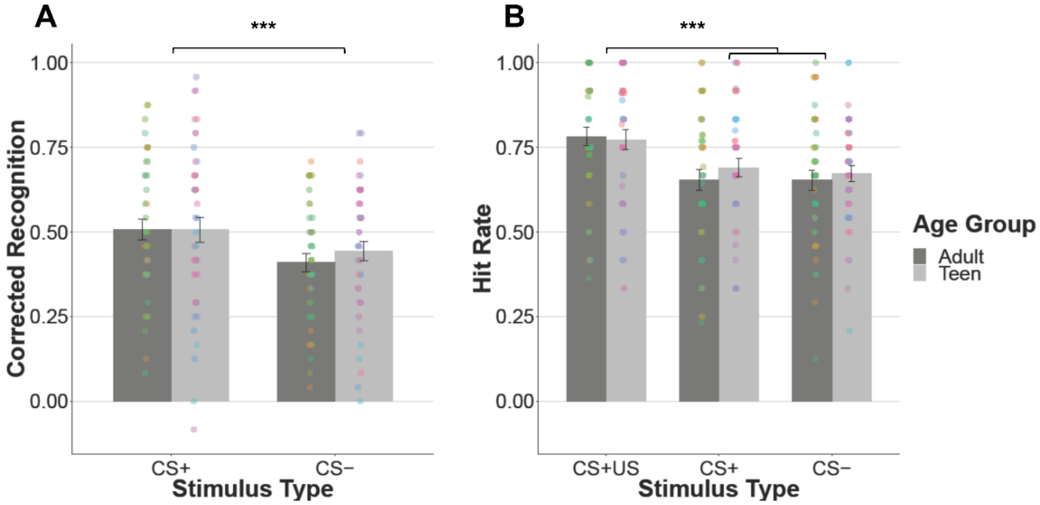


**Figure S1.** Similar effects of aversive learning on recognition memory and skin conductance response across age. Across age, participants show enhanced corrected recognition memory for items from the CS+ versus CS- category (A), driven by better recognition memory for the reinforced items (CS+US) (B). Participants separated by age group for visualization purposes only. Different colored dots represent individual participants. Error bars are s.e.m. *** $p < .001$

We examined hit rate by stimulus type and found a significant effect of stimulus type ($F(2,172) = 16.36$, $p < 0.0001$) and no significant effects of age ($F(1,85) = 0.10$, $p = 0.76$), age by stimulus type interaction ($F(2,172) = 0.57$, $p = 0.57$), or an effect of which category was reinforced ($F(1,85) = 0.05$, $p = 0.82$). Post-hoc t-tests ($\alpha = .0167$, adjusted for multiple comparisons) revealed that the main effect of stimulus type was driven by better memory for the CS+US stimuli relative to the unreinforced CS+ stimuli ($t(173.84) = 3.66$, $p < 0.001$) and the CS- stimuli ($t(173.58) = 4.15$, $p < 0.0001$; Figure S1B). There was no significant difference between memory for the unreinforced CS+ stimuli relative to the CS- stimuli ($t(172.91) = 0.33$, $p = 0.74$). We also examined trial-wise memory accuracy and again found a significant effect of stimulus type ($\chi^2(2) = 49.56$, $p < .0001$) and no significant effects of age ($\chi^2(1) = 0.01$, $p = .90$), age by stimulus type interaction ($\chi^2(2) = 1.99$, $p = .37$), or an effect of which category was reinforced ($\chi^2(1) = 0.28$, $p =.59$). There was a significant primacy effect ($\chi^2(2) = 127.92$, $p < .0001$), such that stimuli presented near the beginning of learning were better remembered than those presented near the end.

*Ordinal Modeling of Recognition Memory Data*

In order to take into account all four potential response options presented to participants in the recognition memory test, we fit an ordinal model in a Bayesian framework using the R package brms (Burkner, 2017; Carpenter et al., 2017; Burkner & Vuorre, 2018). We fit an adjacent category model allowing for category-specific effects of stimulus type. Rather than assuming a predictor has the same effect on all response categories, this model allows a predictor (here, stimulus type) to have different impacts for different response categories (here, 1 = Definitely Old, 2 = Maybe Old, 3 = Maybe New, and 4 = Definitely New) (Burkner & Vuorre, 2018). We included a random intercept for each participant in the model. Age was not included, as we did not find any evidence of age effects in our primary analyses. Credible intervals (CI) that did not overlap with zero are interpreted with a 95% probability of an effect, while those that contain zero are considered small and uncertain effects. We found more high confidence hits (1 = Definitely Old) than low confidence hits (2 = Maybe Old) for CS+US stimuli (b = -0.44, 95%-CI = [-0.58, -0.29]) and CS+ stimuli (b = -0.15, 95%-CI = [-0.30, -0.00]) relative to the CS- stimuli in our main sample of 60 participants (Table S1). We found similar results when including subjects that were excluded for lack of a variable skin-conductance signal (Table S2). These results provide some evidence for generalization of memory facilitation to non-reinforced images in the same category when examining all potential response options, although the effects of memory facilitation for reinforced images remain more robust.

**Table S1.** Summary of regression coefficients for category-specific adjacent category model, n = 60

| | Estimate | l-95% CI | u-95% CI |
|---|---|---|---|
| Intercept[1] | 0.36 | 0.25 | 0.46 |
| Intercept[2] | 0.04 | -0.08 | 0.15 |
| Intercept[3] | 0.42 | 0.29 | 0.55 |
| CS+[1] | -0.15 | -0.30 | -0.00 |
| CS+[2] | 0.09 | -0.08 | 0.27 |
| CS+[3] | 0.00 | -0.19 | 0.20 |
| CS+US[1] | -0.44 | -0.58 | -0.29 |
| CS+US[2] | -0.12 | -0.32 | 0.08 |
| CS+US[3] | 0.05 | -0.17 | 0.28 |

**Table S2.** Summary of regression coefficients for category-specific adjacent category model, n = 88

| | Estimate | l-95% CI | u-95% CI |
|---|---|---|---|
| Intercept[1] | 0.41 | 0.33 | 0.49 |
| Intercept[2] | 0.05 | -0.04 | 0.15 |
| Intercept[3] | 0.39 | 0.29 | 0.29 |
| CS+[1] | -0.13 | -0.25 | -0.01 |
| CS+[2] | 0.02 | -0.12 | 0.18 |
| CS+[3] | 0.09 | -0.07 | 0.25 |
| CS+US[1] | -0.44 | -0.56 | -0.32 |
| CS+US[2] | -0.05 | -0.22 | 0.11 |
| CS+US[3] | 0.02 | -0.17 | 0.20 |

*Recognition Memory-Individual Difference Measure Relationships*

Linear regressions revealed no relationships between CS+US memory bias and age ($F(1,86) = 0.01$, $p = 0.93$) or CS+ memory bias and age ($F(1,86) = 0.71$, $p = 0.40$).

We next examined the relationships between the State Trait Anxiety Index (STAI) state and trait scales and these memory bias measures ($\alpha = .0125$, adjusted for multiple comparisons). We did not find statistically significant relationships between the STAI state measure and either memory bias index (CS+US memory bias, $r(85) = -0.18$, $p = 0.09$; CS+ bias, $r(85) = -0.13$, $p = 0.22$). However, we did observe a negative correlation between the STAI trait and CS+US memory bias ($r(85) = -0.28$, $p = 0.010$) and the STAI trait and CS+ memory bias ($r(85) = -0.33$, $p = 0.002$), such that individuals with lower STAI trait scores showed a stronger CS+US and CS+ memory bias than those with high STAI trait scores. A follow-up analysis ($\alpha = .017$, adjusted for multiple comparisons) was conducted to determine whether these results were due to differences in recognition memory for the CS+US, CS+, or CS- stimuli. We found that neither CS+US hit rate ($r(85) = 0.02$, $p = 0.86$) nor CS+ hit rate ($r(85) = -0.03$, $p = 0.80$) correlated with STAI trait scores. However, we observed a positive correlation between CS- rate and STAI ($r(85) = 0.36$, $p < 0.001$), such that individuals with higher trait anxiety showed better memory for the CS- stimuli. Further analyses ($\alpha = .025$, adjusted for multiple comparisons) revealed that this relationship was statistically significant in the adolescent ($r(42) = 0.43$, $p = 0.003$) and marginally significant in the adult subjects ($r(41) = 0.31$, $p = 0.04$).

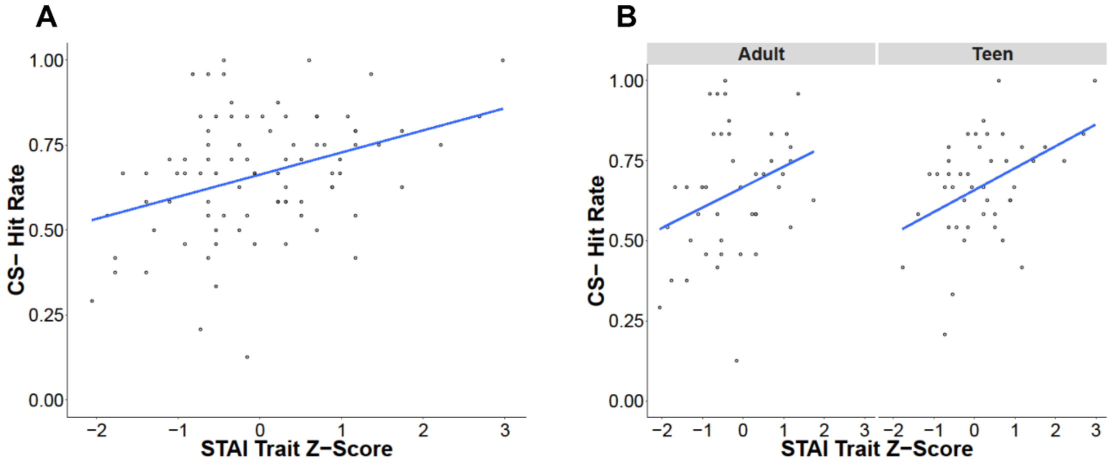


**Figure S2.** Better memory for the stimuli from the unreinforced category is associate with higher trait anxiety. While there was no significant relationship between recognition memory for the CS+ or CS+US stimuli and trait anxiety, there was a positive relationship between memory for the CS- stimuli and trait anxiety (A). Follow-up analysis revealed that this positive relationship was more pronounced in adolescents than adults (B).

We also examined the relationships between Intolerance of Uncertainty Scale (IUS) (Freeston, 1994) and the memory bias measures ($\alpha = .025$, adjusted for multiple comparisons). We did not observe statistically significant correlations between either memory bias index and IUS (CS+US memory bias, $r(85) = -0.14$, $p = 0.20$; CS+ bias, $r(85) = -0.26$, $p = 0.01$). Linear regressions revealed no significant relationships between age and STAI state ($F(1,85) = 1.50$, $p = 0.22$), STAI trait ($F(1,85) = 1.91$, $p = 0.17$), or IUS ($F(1,85) = 0.18$, $p = 0.67$).
